# GABA_A_ and NMDA receptor density alterations and their behavioral correlates in the gestational methylazoxymethanol acetate model for schizophrenia

**DOI:** 10.1101/2021.06.21.449343

**Authors:** Amanda Kiemes, Felipe V. Gomes, Diana Cash, Daniela L. Uliana, Camilla Simmons, Nisha Singh, Anthony C. Vernon, Federico Turkheimer, Cathy Davies, James M. Stone, Anthony A. Grace, Gemma Modinos

## Abstract

Hippocampal hyperactivity driven by GABAergic interneuron deficits and NMDA receptor hypofunction is associated with the hyperdopaminergic state often observed in schizophrenia. Furthermore, previous research in the methylazoxymethanol acetate (MAM) rat model has demonstrated that repeated peripubertal diazepam administration can prevent the emergence of adult hippocampal hyperactivity, dopamine system hyperactivity, and associated psychosis-relevant behaviors. Here, we sought to characterize hippocampal GABA_A_ and NMDA receptors in MAM-treated rats and to elucidate the receptor mechanisms underlying the promising effects of peripubertal diazepam exposure. Quantitative receptor autoradiography was used to measure receptor density in dorsal hippocampus CA1, ventral hippocampus CA1, and in ventral subiculum. Specifically, [^3^H]-Ro15-4513 was used to quantify the density of α5 GABA_A_ receptors (α5GABA_A_R), [^3^H]-flumazenil to quantify α1-3;5GABA_A_R, and [^3^H]-MK801 to quantify NMDA receptors. MAM rats exhibited anxiety and schizophrenia-relevant behaviors as measured by elevated plus maze and amphetamine-induced hyperlocomotion (AIH), although diazepam only partially rescued these behaviors. α5GABA_A_R density was reduced in MAM-treated rats in all hippocampal sub-regions, and negatively correlated with AIH. Ventral hippocampus CA1 α5GABA_A_R density was positively correlated with anxiety-like behavior. Dorsal hippocampus CA1 NMDA receptor density was increased in MAM-treated rats, and positively correlated with AIH. [^3^H]-Flumazenil revealed no significant effects. Finally, we found no significant effect of diazepam treatment on receptor densities, potentially related to the only partial rescue of schizophrenia-relevant phenotypes. Overall, our findings provide first evidence of α5GABA_A_R and NMDA receptor abnormalities in the MAM model, suggesting that more selective pharmacological agents may become a novel therapeutic mechanism in schizophrenia.

## Background

Hippocampal dysfunction has been proposed to underlie the subcortical hyperdopaminergic state commonly associated with psychotic disorders such as schizophrenia [1, 2]. Human *post-mortem* studies revealed reduced hippocampal volume and morphological changes in patients with schizophrenia compared to controls [3, 4]. Consistent with these observations, *in vivo* neuroimaging studies have also documented hippocampal volume reductions [5, 6], and functional abnormalities involving increased hippocampal metabolism, blood flow and activation [7–9]. Moreover, alterations in hippocampal morphology and function have also been observed in individuals at high risk for psychosis [10–13], suggesting that hippocampal dysfunction is already present in psychosis vulnerability states.

Prevailing theories suggest hippocampal hyperactivity in schizophrenia is due to deficits in GABAergic inhibition [1,2,14]. In particular, GABAergic abnormalities in the hippocampus have been found in patient *post-mortem* samples, including lower levels of the GABA synthesizing enzyme GAD67 [15], and a functional loss of parvalbumin-expressing interneurons [16]. Reduced synthesis and release of GABA has been proposed to also lead to a compensatory increase of GABA receptors [17]. Endogenous GABA can act on two different types of receptors, GABA_A_ (GABA_A_R) and GABA_B_ receptors. GABA_A_R are characterized by a wide range of structural diversity and region-specific distribution, due to the large variety of GABA_A_R subunit types [18]. For example, classical benzodiazepines have a high affinity for GABA_A_R at the benzodiazepine binding site (GABA_A_-BZR) at the junction between α1, α2, α3, or α5 subunits and the γ2 subunit, and a low affinity for most other receptor subtypes [19, 20]. Within the hippocampus, α5 subunit-containing GABA_A_Rs (α5GABA_A_R) make up 25% of GABA_A_R in this region [21], but less than 5% of all GABA_A_R in the brain [18]. Additionally, this receptor is critically involved in stress responsivity and sensorimotor gating [18], both commonly disrupted in schizophrenia and high-risk states [22–24]. In *post-mortem* tissue samples of patients with schizophrenia, GABA_A_R were found to be increased in the hippocampus, as measured with quantitative receptor autoradiography using [^3^H]-muscimol [25]. *Post-mortem* studies, however, may be influenced by effects of long-term exposure to medication [26, 27] and illness chronicity on brain tissue as well as technical challenges of tissue condition and preservation. In this context, however, results from *in vivo* neuroimaging studies using positron emission tomography (PET) and ligands that bind to the GABA_A_-BZR ([^123^I]-iomazenil, [^18^F]-fluoroflumazenil, [^11^C]-flumazenil) have been inconsistent [28]. Interestingly, the inverse agonist [^11^C]-Ro15-4513 enables measurement of the α5GABA_A_R more selectively and has also been used in schizophrenia research. From the two existing [^11^C]-Ro15-4513 studies, one including medicated and non-medicated schizophrenia patients found no significant effects [29], while a more recent study reported lower α5GABA_A_R in unmedicated patients only [26].

Preclinical research has provided additional evidence for the relationship between GABAergic abnormalities, hippocampal hyperactivity and the hyperdopaminergic state. A well-validated rat model of relevance for schizophrenia, the methylazoxymethanol acetate (MAM) gestation day (GD) 17 [30] model, introduces a neurodevelopmental insult in the offspring of MAM-treated dams. Their offspring display several behavioral, neuroanatomical and electrophysiological deficits which recapitulate hallmark schizophrenia-relevant features [31]. These include amphetamine-induced hyperlocomotion (AIH) indexing increased striatal dopamine activity [32–34], and anxiety-like behavior in the elevated plus maze (EPM) [33,35,36]. MAM model studies also suggest that hippocampal hyperactivity results from a functional loss of parvalbumin-expressing interneurons [37]. Tonic inhibition provided through α5GABA_A_R provides crucial regulation of glutamatergic pyramidal neuron activity [38]. Importantly, systemic administration of an α5GABA_A_R positive allosteric modulator to MAM-treated adult rats normalized dopamine signaling and reduced the AIH abnormalities [32], and overexpression of hippocampal α5GABA_A_R via viral-mediated gene transfer was shown to also improve schizophrenia-relevant behaviors in the MAM model [39]. With important implications for prophylactic psychiatry, repeated administration of the anxiolytic drug diazepam to MAM rats during the peripubertal period prevented the emergence of parvalbumin-expressing interneuron loss [36], subcortical hyperdopaminergia and elevated stress response [33] in adulthood. Diazepam, a classical benzodiazepine binding to GABA_A_-BZR [20], enhances GABAergic signaling by increasing the affinity of GABA and increasing GABA_A_R channel opening frequency [40], but its wider activity profile may involve acting on both tonic and phasic inhibition within the hippocampus [41]. However, the molecular mechanisms by which diazepam enacts its preventative effects on schizophrenia-relevant pathology in the MAM model is unknown.

The present study aimed to address this issue by using quantitative receptor autoradiography to characterize hippocampal GABAergic and glutamatergic receptor systems in the context of MAM pathophysiology, and the potential modulatory effects of repeated peripubertal diazepam administration on these systems. Specifically, we focused on α5GABA_A_R, GABA_A_R-BZR (α1-3;α5), and NMDA receptor (NMDAR) density. To increase translational relevance, associations between receptor density and behavioral correlates relevant to schizophrenia were examined (i.e. anxiety in the EPM and AIH). Given prior animal research implicating the α5 subunit in the pathophysiology and rescue of schizophrenia-relevant deficits [32,39,42,43], we hypothesized that MAM-treated rats would show reduced α5GABA_A_R binding in the hippocampus, particularly in the ventral portion [44, 45], and that this deficit would be rescued by diazepam treatment. We further hypothesized that unspecific GABA_A_R-BZR would be unaltered in MAM-treated rats but increased by diazepam treatment. Finally, we hypothesized NMDAR density would be increased in MAM-treated rats based on prior human *post-mortem* studies [46], but that it would remain unaffected by diazepam treatment. Because of prior research implicating the CA1 subfield of the ventral hippocampus (vHipp CA1) [17, 47] and the ventral subiculum of the hippocampus (vSub) [44] in the pathogenesis of psychosis, we focused on these regions. Additionally, we explored the CA1 subfield of the dorsal hippocampus (dHipp CA1), given previous evidence demonstrating alterations of GABA_A_R in this region through antipsychotic exposure [27].

### Methods and Materials Animals

All experiments were conducted in accordance with the USPHS’s Guide for the Care and Use of Laboratory Animals and were approved by the Institutional Animal Care and Use Committee of the University of Pittsburgh. Pregnant Sprague-Dawley dams were obtained on GD 15 (Envigo, Indianapolis, IN) and injected with saline (SAL) or MAM (20 mg/kg, intraperitoneal (i.p.); Midwest Research Institute, Kansas City, MO) on GD17. Animals were housed in a 12-hour light/dark cycle (lights on at 7am) in a temperature-(22 ± 1 °C) and humidity-controlled environment.

### Experimental Design

Male pups were weaned on postnatal day (PD) 21 and housed two to three per cage. Animals received diazepam (DZP) or vehicle (VEH) once daily during the peripubertal period (PD 31 – 40). Litters from 7 MAM- and SAL-treated dams each were divided and assigned to the diazepam (SAL:DZP, n=20; MAM:DZP, n=17) and vehicle (SAL:VEH, n=19; MAM:VEH, n=15) group. Once animals reached adulthood (PD 62), they underwent behavioral experiments: (i) elevated plus maze (EPM) to examine anxiety responses, followed by (ii) amphetamine-induced hyperlocomotion (AIH) to test sensitivity to psychostimulants as a proxy for whether MAM exposure had been effective. Subsequently, brains were collected for autoradiography. All animals underwent behavioral testing and autoradiography.

### Oral Administration of Diazepam

Diazepam was administered orally based on a previous study [33]. Briefly, diazepam (5mg/kg, Hospira, INC., Lake Forest, IL) was delivered in wafers (mini Nilla Wafers; Kraft Food) and topped with liquid sugar and sweetened condensed milk (Eagle Brand).

### Elevated Plus Maze (EPM)

The EPM (San Diego Instruments, San Diego, CA) consisted of four 50 cm long and 10 cm wide elevated arms in a cross-like shape positioned 50 cm above the floor. Two opposite arms were enclosed by 40 cm high opaque walls, and the other two were without any walls. A central platform (10 x 10 cm^2^) connected the four arms. Rats were habituated to the testing room for 1 h before the test. For the test, each rat was placed on the central platform, facing an enclosed arm. Behavior was recorded for 5 min. The arena was cleaned between rats. Both the percentage of time spent in the open arms and the percentage of entries into the open arms were taken as measurements of anxiety behavior.

### Amphetamine-Induced Hyperlocomotion (AIH)

Rats were placed into an open-field arena (Coulbourn Instruments, Allentown, PA). Animals’ spontaneous locomotor activity was recorded for 30 min before being injected with D-amphetamine sulfate (0.5 mg/kg, i.p.) after which their locomotor activity was recorded for another 90 min. Locomotor activity was measured via beam breaks and recorded with TruScan software (Coulbourn Instruments). The arena was cleaned between rats.

### Tissue preparation

Rats were anaesthetized with isoflurane (Covetrus, Dublin, OH) and decapitated at PD 69. Brains were carefully removed and flash frozen in isopentane at -40°C. Brains were shipped frozen on dry ice to the BRAIN Centre (Institute of Psychiatry, Psychology & Neuroscience, London, UK), where coronal sections (20 µm thick) were cut in series using a cryostat (Leica CM1950). Sections were mounted onto gelatin coated glass slides and stored at -80°C until used for autoradiography.

### Quantitative receptor autoradiography

Quantitative autoradiography with the radiotracers [^3^H]-Ro15-4513 and [^3^H]-flumazenil was conducted as described previously [27]. To quantify α5GABA_A_R density, we used [^3^H]-Ro15-4513 (Perkin Elmer, NET925250UC). This radiolabeled ligand has a 10- to 15-fold higher affinity to α5GABA_A_R compared to remaining subtypes [48], yielding a high selectivity (60-70%) for the α5 subunit [49] with a smaller portion of selectivity for the α1 subunit [50]. This ligand’s binding pattern highly covaries with the expression of GABRA5, the gene encoding for the α5 subunit [51]. To quantify non-specific binding, we used bretazenil (Sigma, B6434-25MG). Sections were pre-incubated in Tris buffer (50 mM) for 20 min. Slides were subsequently incubated in either 2 nM [^3^H]-Ro15-4513 in Tris buffer for specific binding, or 10 µM bretazenil with 2 nM [^3^H]-Ro15-4513 in Tris buffer for non-specific binding for 60 min. Sections were washed with Tris buffer (2 x 2 min), dipped in distilled water and laid out to dry overnight. All solutions were at room temperature. Dry slides were placed into light-proof cassettes alongside a radioactive [^3^H]-standards slide (American Radiolabelled Chemicals, Inc., USA, ART-123A). A [^3^H]-sensitive film (Amersham Hyperfilm, 28906845) was placed on top of the radioactive slides and exposed for 8 weeks. The film was subsequently developed with a Protex Ecomax film developer (Protec GmbH & Co, Germany).

Similar procedures were used with [^3^H]-flumazenil (Perkin Elmer, NET757001MC), which was used to quantify GABA_A_-BZR binding [52]. Solutions concentrations were 1 nM [^3^H]-flumazenil for total binding, and 10 µM flunitrazepam (Sigma Aldrich, F-907 1ML) with 1 nM [^3^H]-flumazenil for non-specific binding. These were incubated for 60 min and washed in Tris buffer both at 4°C. Film exposure time was 4 weeks long.

To quantify NMDAR, [^3^H]-MK801 (Sigma, M107-25MG) was used with similar procedures as above. Slides were incubated in a 5 nM [^3^H]-MK801 solution for total binding, or in a 5 nM [^3^H]-MK801 solution with the addition of 10 µM MK801 for non-specific binding. Incubation time was 120 min at room temperature. Dried slides were exposed to film for 4 weeks.

### Quantification of receptor binding

Developed films were captured using a Nikon SLR camera. Images of [^3^H]-Ro15-4513 and [^3^H]-flumazenil binding were preprocessed (see supplementary methods). Using MCID software (Imaging Research Inc., 2003), we sampled optical density (OD) values from three primary regions of interests (ROIs) bilaterally (Fig.1): dHipp CA1, vHipp CA1, and vSub. Four secondary ROIs were also sampled (see supplementary methods, Fig. S1). Anatomical regions were defined with the use of Paxinos and Watson’s rat brain atlas [53]. Receptor binding (nCi/mg) was calculated with robust regression interpolation in GraphPad Prism (v9.1.1 for Macintosh, Graphpad Software, La Jolla, CA) using standard curves created from OD measurements of [^3^H]-standards slide for each film. Non-specific binding for [^3^H]-Ro15-4513 and [^3^H]-flumazenil were negligible (see Fig. S2). [^3^H]-MK801 non-specific binding was subtracted from total binding values to calculate [^3^H]-MK801 specific binding.

**Figure 1.**
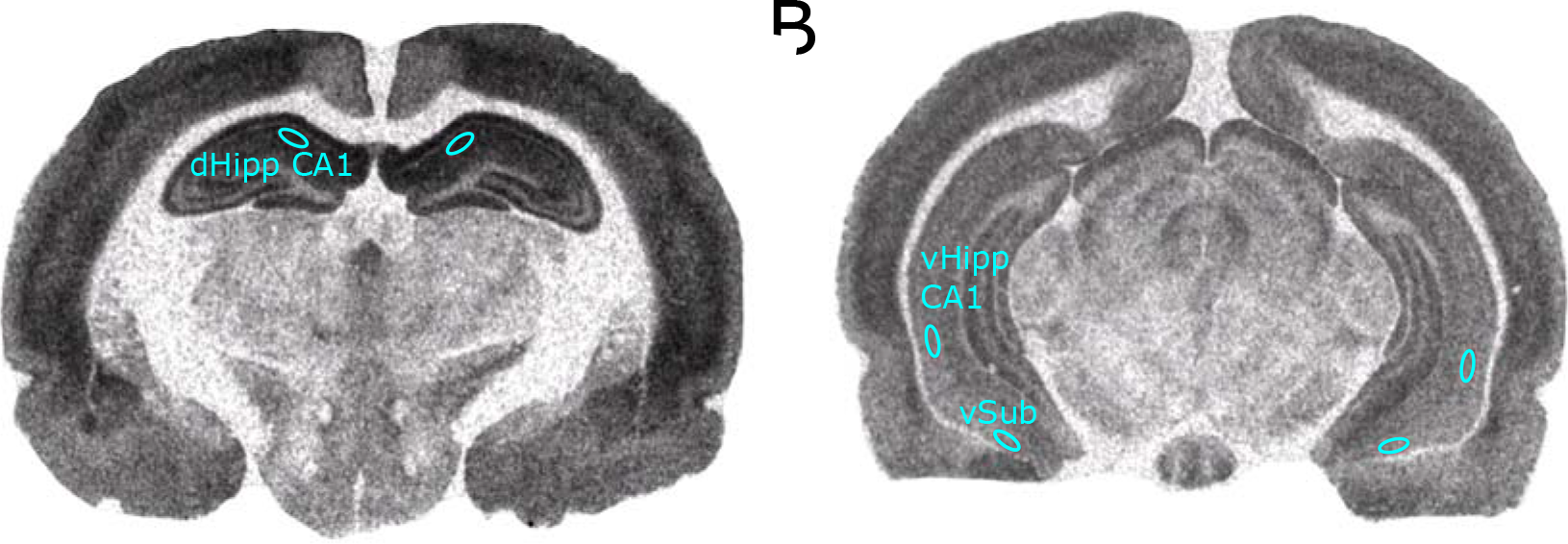
Representative [^3^H]-Ro15-4513 autoradiograph displaying ROI placement. The same ROI placement was used for [^3^H]-flumazenil and [^3^H]-MK801. (A) dorsal hippocampus CA1 (dHipp CA1). (B) ventral hippocampus CA1 (vHipp CA1), ventral subiculum (vSub)

### Statistical analysis

Statistical analysis was performed using GraphPad Prism software (v9.1.1 for Macintosh). Due to failures of the TruScan software, one animal was excluded from analysis for the AIH. Further animals were excluded in the autoradiography data due to failures of the autoradiography protocol: nine animals were excluded for [^3^H]-Ro15-4513 and [^3^H]-Flumazenil and two animals were excluded for [^3^H]-MK801 (see supplementary table S1 for n-values per group). To analyze main effects of group (MAM/SAL) and condition (DZP/VEH), 2-way ANOVAs (AIH total movement; EPM) were utilized. For the AIH time-course and the autoradiography data, 3-way mixed ANOVAs were used with group (MAM/SAL) and condition (DZP/VEH) as between-group factors and time (for AIH) or ROI (for autoradiography data) as within-group factor. ROI interaction effects with group were followed up with 2-way mixed ANOVA (between-subject factor: group; repeated measure: ROI). Pearson’s correlations between behavioral measures and receptor density measures were run where autoradiography data showed significant effects. For these analyses, all rats from different groups were pooled together (see correlational values per group per measure in Supplementary Table 2). *Post hoc* tests were performed where appropriate, corrected using Benjamini-Hochberg method (significance set at *q<0.05)* [54]. The significance threshold was set to *p*<0.05 for all other analyses.

## Results

### MAM-induced Behavioral Phenotypes Were Partially Rescued by Peripubertal Diazepam

Adult MAM rats were previously shown to have a heightened anxiety response in the EPM [33,35,36]. This anxiety-like phenotype was once again confirmed by both measures: MAM-treated rats showed less entries into open arms (main effect of group: F_(1,67)_=5.146, *p=*0.027, η^2^=0.071; Fig. 2A), and less time spent in open arms (main effect of group: F_(1,67)_=5.088, p=0.027, η^2^=0.071; Fig. 2B). Anxiety-like behavior was significantly decreased by peripubertal diazepam treatment in the entries into open arms measure (main effect of condition: F_(1,67)_=7.914, *p*=0.006, η^2^=0.106; Fig. 2A), but failed to reach statistical significance for the time spent in open arms measure (main effect of condition: F_(1,67)_=3.729, *p*=0.058, η^2^=0.053; Fig. 2B). No interaction effects were observed for either measure (*p* > 0.05).

**Figure 2.**
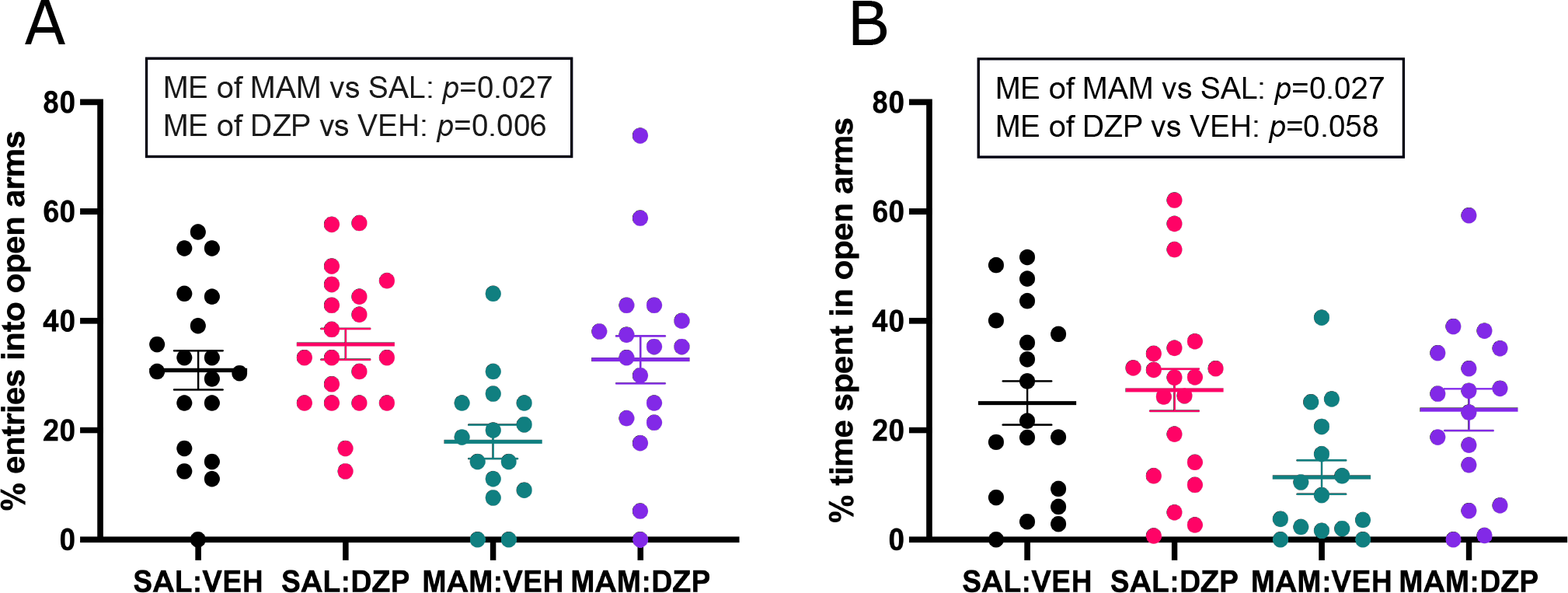
Anxiety-like behavior in the EPM (n=15-20/group). (A) Percentage of entries into open arms. There was a main effect of group (MAM vs SAL) and of condition (DZP vs VEH). (B) Percentage of time spent in the open arms. There is a significant main effect of group (MAM vs SAL), but a main effect of condition (DZP vs VEH) failed to reach significance. Data are displayed as mean ± SEM.

Consistent with previous studies [32–34], MAM-treated adult offspring exhibited increased locomotion in response to amphetamine as compared to SAL-treated adult rats (main effect of group: F_(1,65)_=9.631, *p*=0.003, η^2^=0.156; Fig. 3A). No main effect of condition (F_(1,65)_=2.609, *p*=0.111, η^2^=0.051) or interaction effects (*p*>0.05) were found. Total movement distance post-amphetamine injection further corroborated a significant main effect of group, with increased locomotion in MAM-rats (F_(1,65)_=9.282, *p*=0.003, η^2^=0.125; Fig. 3B), but similarly found no effect of condition (F_(1,65)_=2.163, *p*=0.1462; η^2^=0.032) or interaction effect (F_(1,65)_=0.454, *p*=0.503, η^2^=0.007).

**Figure 3.**
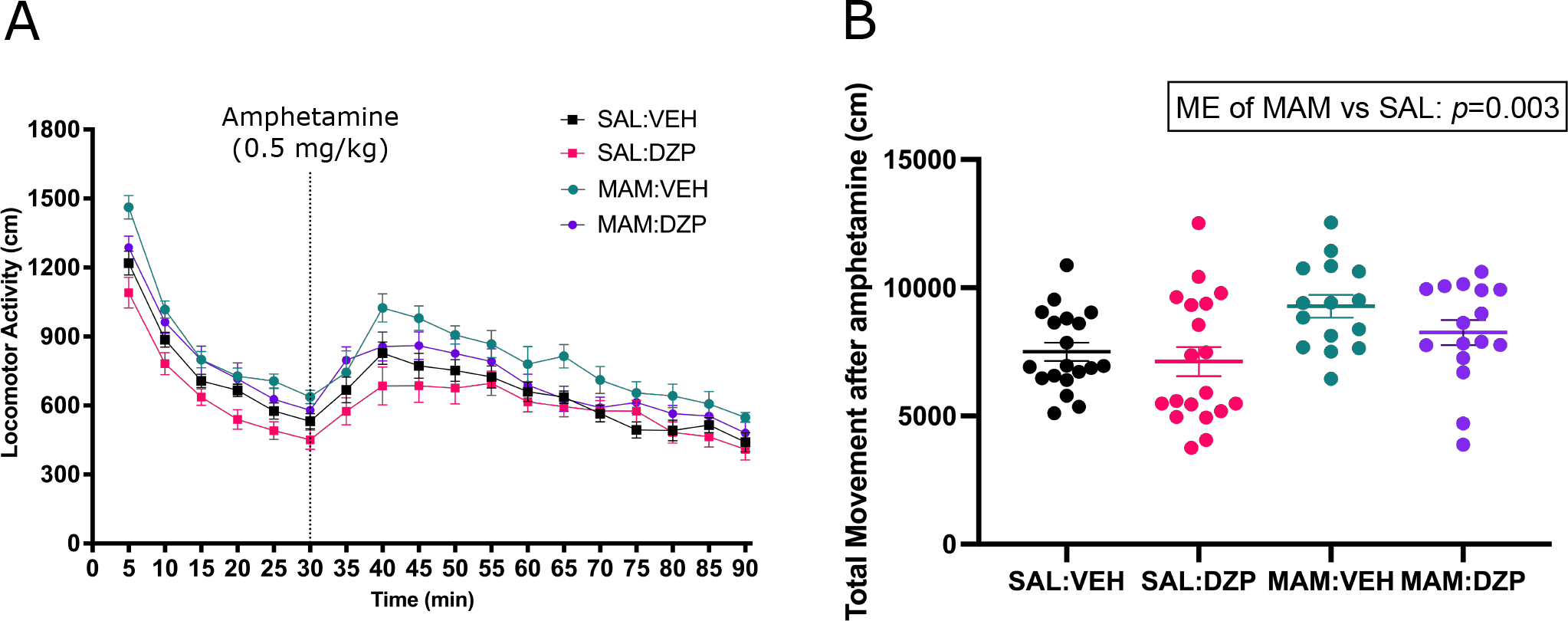
MAM rats showed increased locomotor response to amphetamine compared to controls, which was not prevented by peripubertal diazepam treatment (5mg/kg, oral; daily PD31-40; n=15-20/group). (A) Time course of locomotor activity over 90 minutes. (B) Total movement post amphetamine injection. A main effect of group is observed. Data are displayed as mean ± SEM.

### MAM Treatment Produced Aberrant α5GABA_A_ and NMDA Receptor Density in Adult Rats

α5GABA_A_R density, as indexed by [^3^H]-Ro15-4513 binding, showed a significant main effect of group, with lower binding in MAM rats compared to SAL (F_(1, 58)_=8.410, *p*=0.005, η^2^=0.127; Fig. 4A). There was no significant effect of condition (F_(1,58)_=2.303, *p*=0.135, η^2^=0.038) and no interaction effects were identified (*p*>0.05).

**Figure 4.**
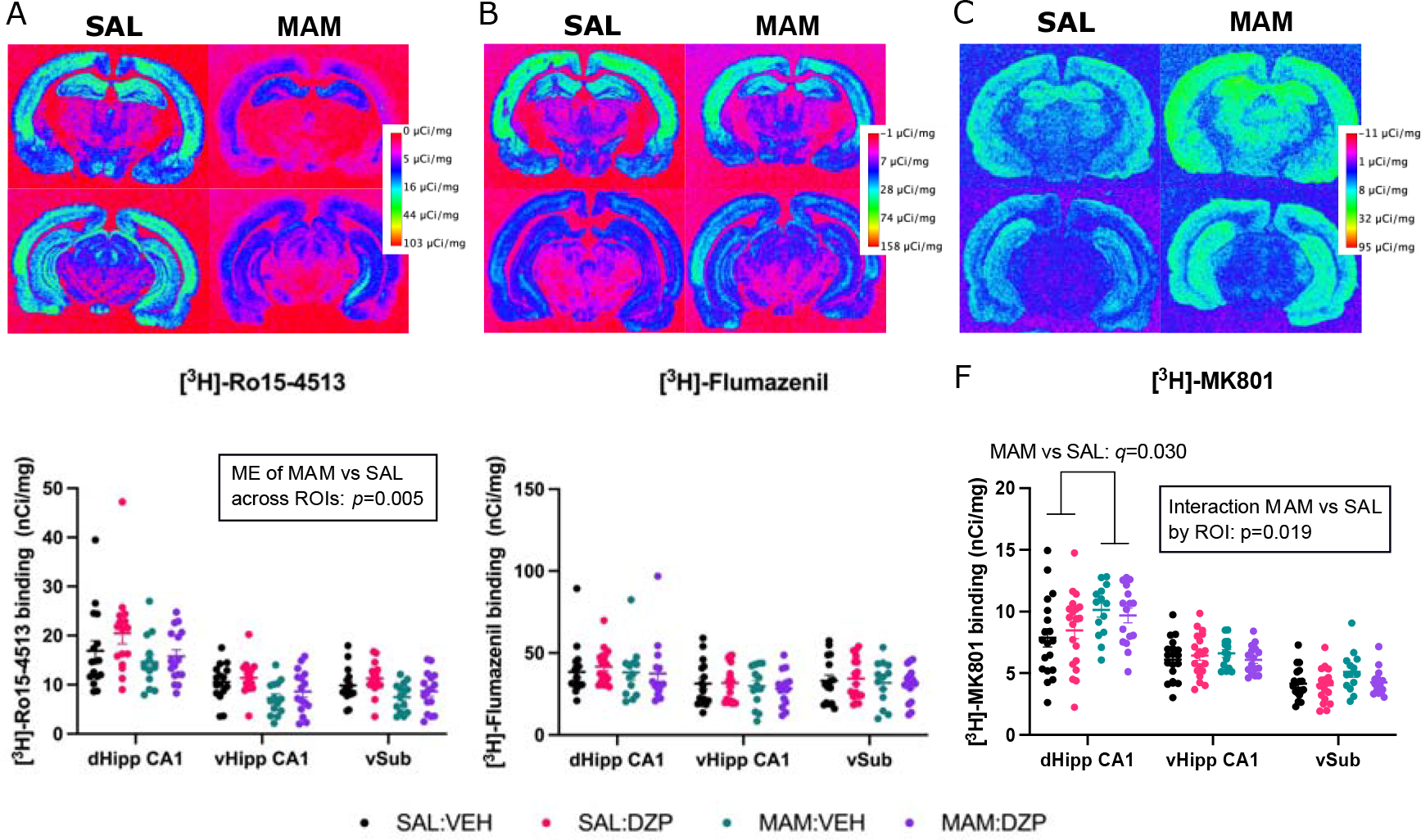
Autoradiography results. Pseudo-color representative autoradiograph of (A) [^3^H]-Ro15-4513 binding (B) [^3^H]-flumazenil binding (C) [^3^H]-MK801 binding. (D) [^3^H]-Ro15-4513 (n=14-16/group) showed a main effect of group. (E) [^3^H]-flumazenil (n=13-18/group) showed no main effects. (D) [^3^H]-MK801 (n=15-20/group) binding showed a main effect of group and a ROI x group interaction. Data are displayed as mean ± SEM.

General GABA_A_-BZR density, as measured with [^3^H]-flumazenil binding, showed no difference between MAM and SAL groups (F_(1,58)_=0.667, *p*=0.417, η^2^=0.011), no effect of condition (diazepam vs vehicle) (F_(1,58)_=0.004, *p*=0.952, η^2^<0.001; Fig. 4B) and no interaction effects (*p*>0.05).

Finally, in terms of [^3^H]-MK801 binding, a main effect of group was observed, with greater binding in MAM compared to SAL (F_(1,65)_=5.483, *p*=0.022, η^2^=0.078; Fig.4C), but no overall effect of condition (F_(1,65)_=0.233, *p*=0.631, η^2^=0.004). Furthermore, an ROI x group interaction effect was found (F_(2,130)_=4.117, *p*=0.019, η^2^=0.060). Follow-up two-way mixed ANOVA, where the SAL:VEH and SAL:DZP groups were combined into one SAL group and the MAM:VEH and MAM:DZP groups into one MAM group, reflected that the effect of group was only significant within the dHipp CA1 region (greater [^3^H]-MK801 binding in MAM vs SAL, *p*=0.001, *q*=0.030). The above autoradiographic analyses were repeated with secondary ROIs included and are presented in the supplementary results.

### α5GABA_A_R and NMDAR density were differentially correlated with schizophrenia-relevant behaviors

Correlation between behavioral measures (AIH total movement, EPM time spent in open arms, EPM entries into open arms) and those receptor density measures that showed a significant group effect ([^3^H]-Ro15-3415 dHipp CA1, vHipp CA1, and vSub; [^3^H]-MK801 dHipp CA1) revealed that all correlations with AIH total movement were significant (Fig.5). Lower [^3^H]-Ro15-4513 receptor binding was associated with greater locomotor response to amphetamine (dHipp CA1: *r*=-0.316, *p*=0.013, *q*=0.013, Fig. 5A; vHipp CA1: *r*=-0.366, *p*=0.004, *q*=0.006, Fig. 5B; vSub: *r*=-0.401, *p*=0.001, *q*=0.003, Fig. 5C). Meanwhile, higher [^3^H]-MK801 receptor binding in the dHipp CA1 was associated with higher locomotor response to amphetamine (*r*=0.318, *p*=0.007, Fig. 5D). In terms of EPM measures, vHipp CA1 [^3^H]-Ro15-4513 binding was significantly positively associated (*r*=0.263, *p*=0.039, Fig. 5F) with time spent in open arms, but positive association with entries into open arms missed significance (*r*=0.240, *p*=0.061, Fig. 5E). No other significant correlations with EPM measures were found. Correlations were also run separately per group and are presented in the supplementary results (Table S3).

**Figure 5.**
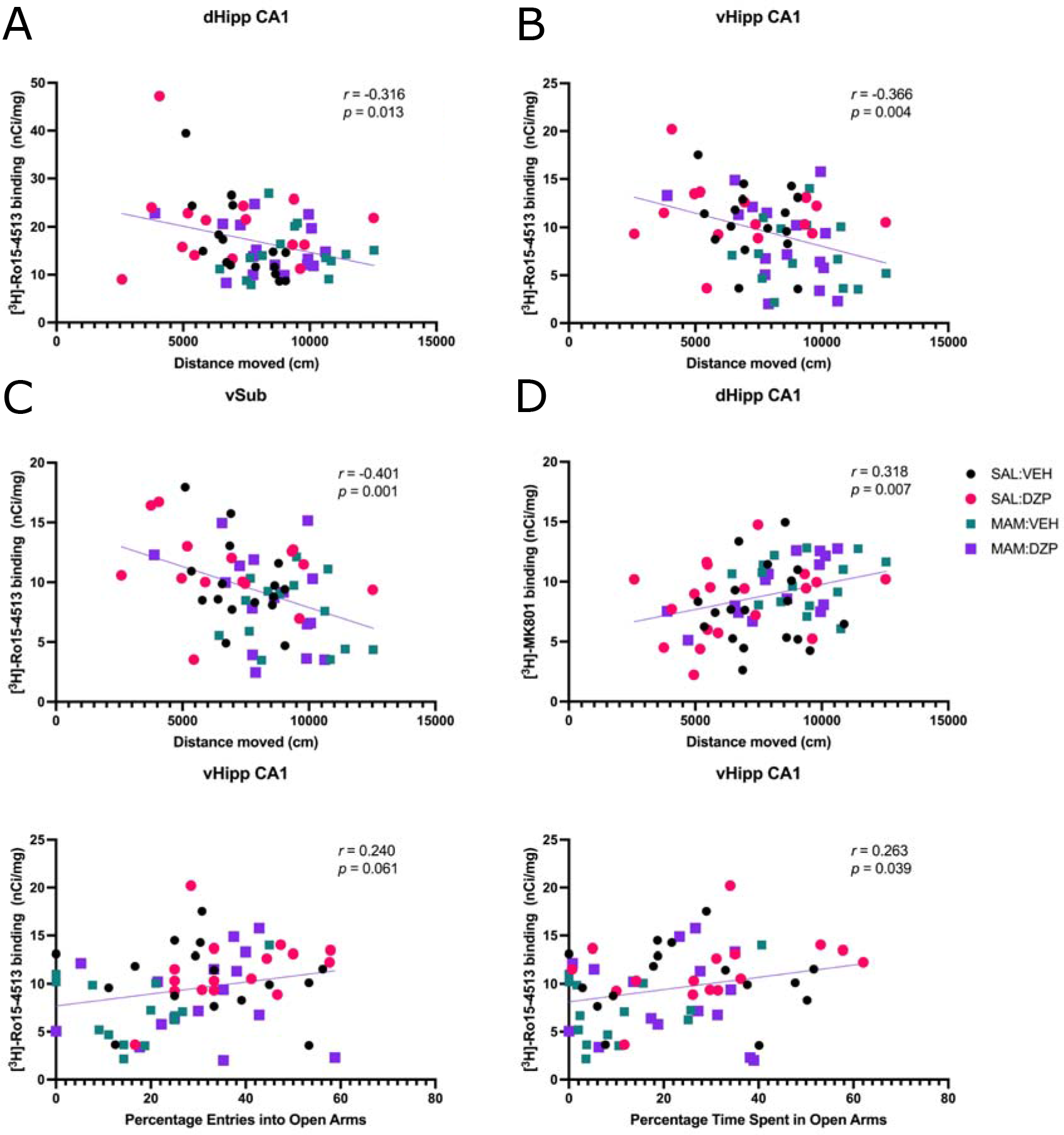
Significant autoradiography x behavior correlations for pooled groups. (A-C) [^3^H]-Ro15-4513 binding was inversely correlated with amphetamine-induced hyperlocomotion. (D) [^3^H]-MK801 binding was positively correlated with amphetamine-induced hyperlocomotion. (E-F) [^3^H]-Ro15-4513 binding was positively correlated with time spent in open arms, but correlation failed to reach significance with entries into open arms of the EPM.

## Discussion

In this study, we sought to characterize GABAergic and glutamatergic receptor abnormalities in hippocampus using a well-validated neurodevelopmental model relevant to schizophrenia, and their potential modulation by peripubertal diazepam treatment. Our main finding was that MAM-treated rats displayed lower α5GABA_A_R density in key hippocampal sub-regions, indexed by lower [^3^H]-Ro15-4513 binding. This reduction appears to be specific to α5GABA_A_R, based on [^3^H]-Ro15-4513’s higher affinity to this receptor, and the lack of significant effects in non-specific GABA_A_R-BZR density as measured by [^3^H]-flumazenil. Furthermore, MAM treatment was associated with greater NMDAR density, specific to the dorsal hippocampus CA1 subfield. Importantly, these receptor abnormalities were linked to schizophrenia-relevant behavioral phenotypes, as both the α5GABA_A_R decrease and the NMDAR increase were associated with AIH. While we had predicted that peripubertal diazepam treatment would prevent the emergence of schizophrenia-relevant behaviors and normalize receptor density abnormalities identified by autoradiography, we observed only a partial behavioral rescue (i.e. anxiety-like behavior was rescued, but AIH was not) of the phenotypes and no treatment effects of diazepam on receptor densities.

Lower α5GABA_A_R density in the MAM model is consistent with our hypothesis, based on previous literature suggesting a key role for this subunit in correcting several aspects of the MAM-related phenotype, including dopamine-system hyperactivity [32, 39]. α5GABA_A_R binding in the hippocampus has been described in unmedicated schizophrenia patients by *in vivo* PET imaging using the same ligand [^11^C]-Ro15-4513 [26]. Our findings thus provide first evidence that a specific deficit in this receptor subtype is involved in MAM pathophysiology relevant to schizophrenia, using a translational imaging measure, and building on previous evidence that selectively targeting this subunit could potentially compensate for schizophrenia pathology [32]. In terms of mechanisms, lower α5GABA_A_R function in vHipp CA1 and vSub may underlie hippocampal hyperactivity in schizophrenia through disinhibition of glutamatergic pyramidal cell activity [32]. Neuroimaging studies in patients with schizophrenia and high-risk individuals have documented elevated cerebral blood volume/flow specifically in the CA1 subfield [13,55,56]. Moreover, elevated cerebral blood volume correlated with positive symptom severity [13, 56], consistent with the inverse relationship we observed between hippocampal α5GABA_A_R density and locomotor response to amphetamine, as α5GABA_A_R regulate tonic inhibition. Interestingly, anxiety-like behavior was exclusively associated with α5GABA_A_R in the vHipp CA1, as no associations were found with any of the other sub-regions or receptor types. Human and animal studies have suggested that hippocampal dysfunction originates in CA1 and spreads to the subiculum, from which the hippocampus then dysregulates ventral tegmental area dopamine neurons [9,47,57]. These findings may suggest that the ventral CA1 may be most intimately linked to psychosis pathophysiology and to the increased stress responsivity thought to be a risk factor for transition [58].

Within the dHipp CA1, we observed lower α5GABA_A_R and increased NMDAR. Implications of lower α5GABA_A_R in the dHipp CA1 remain unclear. Previous research by our group identified a dose-dependent change in α5GABA_A_R density in this region in response to chronic haloperidol exposure [27]. Hence, a possible explanation may relate to a disease-driven receptor abnormality in this region, which may explain the receptor increases in response to haloperidol [27] and provide another mechanism of action for antipsychotics. In terms of NMDAR, the observed increase in dHipp CA1 aligns with a prior autoradiographic study of human *post-mortem* brain tissue from patients with schizophrenia [46]. Specifically, an overall increase of NMDAR in multiple regions including the hippocampus was found; however, only receptors in the putamen reached statistical significance. Kornhuber et al. [46] speculated that this increase may be partially due to effects of antipsychotic exposure; however, our findings suggest that this may not be the case. Due to the putative hypofunction of this receptor in the pathophysiology of schizophrenia [17], increased expression may be a compensatory mechanism that develops alongside schizophrenia progression. The negative correlation of α5GABA_A_R and the positive correlation of NMDAR with AIH would suggest that these receptors are moderately affiliated with behavioral abnormalities relevant to schizophrenia. However, unlike α5GABA_A_R in vHipp CA1, dHipp CA1 receptors were not related to stress measures (EPM). Given that the hippocampus is a functionally segregated structure, with vHipp (anterior in primates) playing a role in emotion and stress while the dHipp (posterior in primates) is involved with information processing [59], future studies including behavioral tasks dependent on dHipp, such as spatial memory and context discrimination [59, 60], will expand on our findings.

In terms of [^3^H]-flumazenil, we found no effects in GABA-BZR density in MAM-treated animals. [^3^H]-flumazenil was used as a positive control to test the specificity of [^3^H]-Ro15-4513 to α5GABA_A_R. In combination with only partial evidence supporting a diazepam effect in the behavioral measures (i.e. only in EPM measures, not in AIH), these observations may reflect that repeated diazepam treatment failed to fully recover the model in our study, in contrast with previous work [33, 36]. In humans, an elevated anxiety response to stress is present in high-risk individuals and associated with the subsequent transition to psychosis [58]. Peripubertal diazepam in the MAM model is thought to prevent schizophrenia-like behavior by attenuating this heightened stress response [33, 36]. However, in humans, on occasion an uncommon (in less than 1% of patients) paradoxical reaction to benzodiazepines occurs: rather than displaying signs of sedation they become agitated, excited, and engage in emotional release and excessive movement [61]. Albeit rare, these reactions have been putatively linked to psychiatric disorders such as bipolar disorder and schizophrenia [61], but the mechanisms still remain elusive. Such reactions illustrate that diazepam acts on a functionally diverse system of feedforward and feedback connections, and synaptic and extrasynaptic receptors [18], and thus the effects of diazepam may not always be predictable.

There are some limitations to our study that should be noted. Firstly, we used AIH as an assay to index schizophrenia-like behavior; however, the analogousness to associative striatum hyperdopaminergia in humans has been debated [62–64]. AIH is rather thought to reflect ventromedial limbic striatal dopamine action and thus cannot be taken as a direct analogue of positive symptoms in animal models. Nonetheless, this behavioral assay is commonly used to study the efficacy of antipsychotic drugs [63]. Future studies using behavioral tests assessing salience attribution and selective attention, which are more directly related to associative striatum dopamine release [64], are warranted. Secondly, autoradiography was only performed at adulthood, as imaging of autoradiographs required termination of the animal and as such cannot be performed longitudinally in the same animals. Our experimental design allowed measuring behavior, drug effects and receptor densities in the same animals. However, this design does not allow tracking neurodevelopmental changes in α5GABA_A_R and NMDAR densities and their potential role MAM pathophysiology. Further *in vivo* studies using these tracers in the context of PET imaging will enable mapping of the trajectory of receptor abnormalities identified in our study.

In summary, the present findings implicate α5GABA_A_R abnormalities in schizophrenia-relevant pathophysiology, and provide new empirical support to the notion that the development of pharmacological agents with selectivity for hippocampal α5GABA_A_R may be a promising new therapeutic target to prevent schizophrenia-related deficits. As our report is, to our knowledge, the first to image GABA_A_R and NMDAR in the MAM model with comparable measures that can be used in humans, future translational studies imaging these receptor subunits in early psychosis are warranted to inform whether clinical interventions targeting this pathway may have the potential to prevent or delay the development of psychosis in vulnerable individuals.

### Funding and Disclosures

AAG received consultant fees from Lundbeck, Pfizer, Otsuka, Asubio, Autofony, Alkermes, Concert and Janssen, and is on the advisory board for Alkermes, Newron and Takeda. In the last 3 years, JMS has been principal investigator or sub-investigator on studies sponsored by Takeda, Janssen and Lundbeck Plc. He has attended an Investigators’ meeting run by Allergan Plc. All other authors report no potential conflicts of interest.

This work was part supported by a Sir Henry Dale Fellowship jointly funded by the Wellcome Trust and the Royal Society to GM (202397/Z/16/Z). This work was in part supported by National Institutes of Health Grant No. MH57440 to AAG. FVG receives a Sao Paulo Research Foundation Young Investigator grant (2018/17597-3).

Data from a subset of animals used in this paper had been previously analysed and published in conference abstracts (https://doi.org/10.1093/schbul/sbaa029.740; https://doi.org/10.1016/j.euroneuro.2021.01.014). A version of this manuscript can be found on the bioRxiv pre-print server (https://doi.org/10.1101/2021.06.21.449343).

## Author Contributions

GM and AAG developed the study concept and experimental design; AK, FVG, CD, DLU, CS and NS were involved in conducting the experiment and data collection; AK, FVG and DLU analyzed the data; all authors helped to interpret the data; AK wrote the manuscript with input from all authors. All authors have approved the manuscript submitted for publication.

## Notes

### Summary of Updates

Title was revised to fit new formatting; Abstract was updated to clarify objectives of study; figures were revised to fit new formatting; disclosures have been updated and expanded; Table was moved to supplemental files

## References

1. Tamminga CA, Stan AD, Wagner AD. The Hippocampal Formation in Schizophrenia. American Journal of Psychiatry. 2010;167(10):1178–93.

2. Heckers S, Konradi C. GABAergic mechanisms of hippocampal hyperactivity in schizophrenia. Schizophrenia Research. 2015;167(1):4–11.

3. Rimol LM, Hartberg CB, Nesvåg R, Fennema-Notestine C, Hagler DJ, Pung CJ, et al. Cortical Thickness and Subcortical Volumes in Schizophrenia and Bipolar Disorder. Biological Psychiatry. 2010;68(1):41–50.

4. Dwork AJ. Postmortem Studies of the Hippocampal Formation in Schizophrenia. Schizophrenia Bulletin. 1997;23(3):385–402.

5. Lawrie SM, Abukmeil SS. Brain abnormality in schizophrenia: A systematic and quantitative review of volumetric magnetic resonance imaging studies. British Journal of Psychiatry. 1998;172(2):110–20.

6. Honea R, Crow TJ, Passingham D, Mackay CE. Regional deficits in brain volume in schizophrenia: a meta-analysis of voxel-based morphometry studies. Am J Psychiatry. 2005;162(12):2233–45.

7. Malaspina D, Harkavy-Friedman J, Corcoran C, Mujica-Parodi L, Printz D, Gorman JM, et al. Resting neural activity distinguishes subgroups of schizophrenia patients. Biol Psychiatry. 2004;56(12):931–7.

8. Kühn S, Gallinat J. Resting-state brain activity in schizophrenia and major depression: a quantitative meta-analysis. Schizophr Bull. 2013;39(2):358–65.

9. Lieberman JA, Girgis RR, Brucato G, Moore H, Provenzano F, Kegeles L, et al. Hippocampal dysfunction in the pathophysiology of schizophrenia: a selective review and hypothesis for early detection and intervention. Molecular Psychiatry. 2018;23(8):1764–72.

10. Pantelis C, Velakoulis D, McGorry PD, Wood SJ, Suckling J, Phillips LJ, et al. Neuroanatomical abnormalities before and after onset of psychosis: a cross-sectional and longitudinal MRI comparison. The Lancet. 2003;361(9354):281–88.

11. Schobel SA, Kelly MA, Corcoran CM, Van Heertum K, Seckinger R, Goetz R, et al. Anterior hippocampal and orbitofrontal cortical structural brain abnormalities in association with cognitive deficits in schizophrenia. Schizophrenia research. 2009;114(1-3):110–18.

12. Allen P, Azis M, Modinos G, Bossong MG, Bonoldi I, Samson C, et al. Increased Resting Hippocampal and Basal Ganglia Perfusion in People at Ultra High Risk for Psychosis: Replication in a Second Cohort. Schizophrenia Bulletin. 2017;44(6):1323–31.

13. Schobel SA, Lewandowski NM, Corcoran CM, Moore H, Brown T, Malaspina D, et al. Differential targeting of the CA1 subfield of the hippocampal formation by schizophrenia and related psychotic disorders. Archives of general psychiatry. 2009;66(9):938–46.

14. Grace AA, Gomes FV. The Circuitry of Dopamine System Regulation and its Disruption in Schizophrenia: Insights Into Treatment and Prevention. Schizophrenia Bulletin. 2019;45(1):148–57.

15. Benes FM, Lim B, Matzilevich D, Walsh JP, Subburaju S, Minns M. Regulation of the GABA cell phenotype in hippocampus of schizophrenics and bipolars. Proceedings of the National Academy of Sciences. 2007;104(24):10164–69.

16. Zhang ZJ, Reynolds GP. A selective decrease in the relative density of parvalbumin-immunoreactive neurons in the hippocampus in schizophrenia. Schizophrenia research. 2002;55(1-2):1–10.

17. Lisman JE, Coyle JT, Green RW, Javitt DC, Benes FM, Heckers S, et al. Circuit-based framework for understanding neurotransmitter and risk gene interactions in schizophrenia. Trends in Neurosciences. 2008;31(5):234–42.

18. Engin E, Benham RS, Rudolph U. An Emerging Circuit Pharmacology of GABA(A) Receptors. Trends Pharmacol Sci. 2018;39(8):710–32.

19. Pym LJ, Cook SM, Rosahl T, McKernan RM, Atack JR. Selective labelling of diazepam-insensitive GABAA receptors in vivo using [3H]Ro 15-4513. Br J Pharmacol. 2005;146(6):817–25.

20. Sigel E, Ernst M. The Benzodiazepine Binding Sites of GABA(A) Receptors. Trends Pharmacol Sci. 2018;39(7):659–71.

21. Sur C, Fresu L, Howell O, Mckernan R, Atack J. Autoradiographic localization of alpha5 subunit-containing GABAA receptors in rat brain. Brain research. 1999;822 1-2:265–70.

22. San-Martin R, Castro LA, Menezes PR, Fraga FJ, Simões PW, Salum C. Meta-Analysis of Sensorimotor Gating Deficits in Patients With Schizophrenia Evaluated by Prepulse Inhibition Test. Schizophrenia Bulletin. 2020;46(6):1482–97.

23. Walker E, Mittal V, Tessner K. Stress and the Hypothalamic Pituitary Adrenal Axis in the Developmental Course of Schizophrenia. Annual Review of Clinical Psychology. 2008;4(1):189–216.

24. Winton-Brown T, Kumari V, Windler F, Moscoso A, Stone J, Kapur S, et al. Sensorimotor gating, cannabis use and the risk of psychosis. Schizophrenia Research. 2015;164(1):21–27.

25. Benes FM, Khan Y, Vincent SL, Wickramasinghe R. Differences in the subregional and cellular distribution of GABAA receptor binding in the hippocampal formation of schizophrenic brain. Synapse. 1996;22(4):338–49.

26. Marques TR, Ashok AH, Angelescu I, Borgan F, Myers J, Lingford-Hughes A, et al. GABA-A receptor differences in schizophrenia: a positron emission tomography study using [11C]Ro154513. Molecular Psychiatry. 2020.

27. Peris-Yague A, Kiemes A, Cash D, Cotel M-C, Singh N, Vernon AC, et al. Region-specific and dose-specific effects of chronic haloperidol exposure on [3H]-flumazenil and [3H]-Ro15-4513 GABAA receptor binding sites in the rat brain. European Neuropsychopharmacology. 2020;41:106–17.

28. Egerton A, Modinos G, Ferrera D, McGuire P. Neuroimaging studies of GABA in schizophrenia: a systematic review with meta-analysis. Translational Psychiatry. 2017;7(6):e1147–e47.

29. Asai Y, Takano A, Ito H, Okubo Y, Matsuura M, Otsuka A, et al. GABAA/Benzodiazepine receptor binding in patients with schizophrenia using [11C]Ro15-4513, a radioligand with relatively high affinity for α5 subunit. Schizophrenia Research. 2008;99(1):333–40.

30. Lodge DJ. The MAM rodent model of schizophrenia. Current protocols in neuroscience. 2013;63(1):9.43.1–9.43.7.

31. Lodge DJ, Grace AA. Gestational methylazoxymethanol acetate administration: a developmental disruption model of schizophrenia. Behav Brain Res. 2009;204(2):306–12.

32. Gill KM, Lodge DJ, Cook JM, Aras S, Grace AA. A Novel α5GABAAR-Positive Allosteric Modulator Reverses Hyperactivation of the Dopamine System in the MAM Model of Schizophrenia. Neuropsychopharmacology. 2011;36(9):1903–11.

33. Du Y, Grace AA. Peripubertal Diazepam Administration Prevents the Emergence of Dopamine System Hyperresponsivity in the MAM Developmental Disruption Model of Schizophrenia. Neuropsychopharmacology. 2013;38(10):1881–88.

34. Flagstad P, Mørk A, Glenthøj BY, van Beek J, Michael-Titus AT, Didriksen M. Disruption of neurogenesis on gestational day 17 in the rat causes behavioral changes relevant to positive and negative schizophrenia symptoms and alters amphetamine-induced dopamine release in nucleus accumbens. Neuropsychopharmacology. 2004;29(11):2052–64.

35. Zhu X, Grace AA. Prepubertal Environmental Enrichment Prevents Dopamine Dysregulation and Hippocampal Hyperactivity in MAM Schizophrenia Model Rats. Biological Psychiatry. 2021;89(3):298–307.

36. Du Y, Grace AA. Amygdala Hyperactivity in MAM Model of Schizophrenia is Normalized by Peripubertal Diazepam Administration. Neuropsychopharmacology. 2016;41(10):2455–62.

37. Lodge DJ, Behrens MM, Grace AA. A Loss of Parvalbumin-Containing Interneurons Is Associated with Diminished Oscillatory Activity in an Animal Model of Schizophrenia. The Journal of Neuroscience. 2009;29(8):2344–54.

38. Semyanov A, Walker MC, Kullmann DM, Silver RA. Tonically active GABAA receptors: modulating gain and maintaining the tone. Trends in neurosciences. 2004;27(5):262–69.

39. Donegan JJ, Boley AM, Yamaguchi J, Toney GM, Lodge DJ. Modulation of extrasynaptic GABA(A) alpha 5 receptors in the ventral hippocampus normalizes physiological and behavioral deficits in a circuit specific manner. Nat Commun. 2019;10(1):2819.

40. Uusi-Oukari M, Korpi ER. Regulation of GABAA Receptor Subunit Expression by Pharmacological Agents. Pharmacological Reviews. 2010;62(1):97–135.

41. Farrant M, Nusser Z. Variations on an inhibitory theme: phasic and tonic activation of GABAA receptors. Nature Reviews Neuroscience. 2005;6(3):215–29.

42. Gerdjikov TV, Rudolph U, Keist R, Möhler H, Feldon J, Yee BK. Hippocampal α5 subunit-containing GABAA receptors are involved in the development of the latent inhibition effect. Neurobiology of Learning and Memory. 2008;89(2):87–94.

43. Hauser J, Rudolph U, Keist R, Möhler H, Feldon J, Yee BK. Hippocampal α5 subunit-containing GABAA receptors modulate the expression of prepulse inhibition. Molecular Psychiatry. 2005;10(2):201–07.

44. Grace AA. Dopamine System Dysregulation by the Ventral Subiculum as the Common Pathophysiological Basis for Schizophrenia Psychosis, Psychostimulant Abuse, and Stress. Neurotoxicity Research. 2010;18(3):367–76.

45. Lodge DJ, Grace AA. Aberrant Hippocampal Activity Underlies the Dopamine Dysregulation in an Animal Model of Schizophrenia. The Journal of Neuroscience. 2007;27(42):11424–30.

46. Kornhuber J, Mack-Burkhardt F, Riederer P, Hebenstreit G, Reynolds G, Andrews H, et al. (3 H] MK-801 binding sites in postmortem brain regions of schizophrenic patients. Journal of Neural Transmission. 1989;77(2-3):231–36.

47. Schobel SA, Chaudhury NH, Khan UA, Paniagua B, Styner MA, Asllani I, et al. Imaging patients with psychosis and a mouse model establishes a spreading pattern of hippocampal dysfunction and implicates glutamate as a driver. Neuron. 2013;78(1):81–93.

48. Hadingham KL, Wingrove P, Le Bourdelles B, Palmer KJ, Ragan CI, Whiting PJ. Cloning of cDNA sequences encoding human alpha 2 and alpha 3 gamma-aminobutyric acidA receptor subunits and characterization of the benzodiazepine pharmacology of recombinant alpha 1-, alpha 2-, alpha 3-, and alpha 5-containing human gamma-aminobutyric acidA receptors. Mol Pharmacol. 1993;43(6):970–5.

49. Myers JF, Comley RA, Gunn RN. Quantification of [11C]Ro15-4513 GABAAα5 specific binding and regional selectivity in humans. Journal of Cerebral Blood Flow & Metabolism. 2017;37(6):2137–48.

50. Myers JF, Rosso L, Watson BJ, Wilson SJ, Kalk NJ, Clementi N, et al. Characterisation of the Contribution of the GABA-Benzodiazepine α1 Receptor Subtype to [11C]Ro15-4513 PET Images. Journal of Cerebral Blood Flow & Metabolism. 2012;32(4):731–44.

51. Lukow P, Martins D, Veronese M, Vernon A, McGuire P, Turkheimer F, et al. Cellular and molecular signatures of

52. *in vivo* GABAergic neurotransmission in the human brain. bioRxiv. 2021:2021.06.17.448812.

53. Sieghart W. Structure, pharmacology, and function of GABAA receptor subtypes. Adv Pharmacol. 2006;54:231–63.

54. Paxinos G, Watson C. The rat brain in stereotaxic coordinates. 5 ed. Elsevier Academic Press; 2004.

55. Benjamini Y, Hochberg Y. Controlling the false discovery rate: a practical and powerful approach to multiple testing. Journal of the Royal statistical society: series B (Methodological). 1995;57(1):289–300.

56. Talati P, Rane S, Kose S, Blackford JU, Gore J, Donahue MJ, et al. Increased hippocampal CA1 cerebral blood volume in schizophrenia. NeuroImage: Clinical. 2014;5:359–64.

57. Modinos G, Richter A, Egerton A, Bonoldi I, Azis M, Antoniades M, et al. Interactions between hippocampal activity and striatal dopamine in people at clinical high risk for psychosis: relationship to adverse outcomes. Neuropsychopharmacology. 2021;46(8):1468–74.

58. Sonnenschein SF, Gomes FV, Grace AA. Dysregulation of Midbrain Dopamine System and the Pathophysiology of Schizophrenia. Frontiers in Psychiatry. 2020;11(613).

59. Owens DG, Miller P, Lawrie SM, Johnstone EC. Pathogenesis of schizophrenia: a psychopathological perspective. Br J Psychiatry. 2005;186:386–93.

60. Fanselow MS, Dong H-W. Are the Dorsal and Ventral Hippocampus Functionally Distinct Structures? Neuron. 2010;65(1):7–19.

61. Frankland PW, Cestari V, Filipkowski RK, McDonald RJ, Silva AJ. The dorsal hippocampus is essential for context discrimination but not for contextual conditioning. Behavioral neuroscience. 1998;112(4):863.

62. Mancuso CE, Tanzi MG, Gabay M. Paradoxical Reactions to Benzodiazepines: Literature Review and Treatment Options. Pharmacotherapy: The Journal of Human Pharmacology and Drug Therapy. 2004;24(9):1177–85.

63. Carpenter WT, Koenig JI. The evolution of drug development in schizophrenia: past issues and future opportunities. Neuropsychopharmacology. 2008;33(9):2061–79.

64. Nestler EJ, Hyman SE. Animal models of neuropsychiatric disorders. Nature neuroscience. 2010;13(10):1161.

65. Sonnenschein SF, Grace AA. Insights on current and novel antipsychotic mechanisms from the MAM model of schizophrenia. Neuropharmacology. 2020;163:107632.

