## Supplementary Information for "GABA_A_ and NMDA receptor density alterations and their behavioral correlates in the gestational methylazoxymethanol acetate model for schizophrenia"

### Supplementary Methods

#### Autoradiography development and image processing

Developed autoradiographic films were placed on top of a light box (Northern Lights, USA). Images were captured using an AF-S Micro NIKKOR 60 mm F2.82G ED lens. Uniform lighting conditions were kept during image capture. Images were preprocessed on Adobe Photoshop (Adobe Photoshop CC for Macintosh, https://www.adobe.com). Pre-processing steps were conversion to gray followed by contrast normalization. Normalization was achieved by using the 'Levels' tool and selecting the film background as white and the maximum standard used (63.1 nCi/mg) as black. Automatic batch processing was then completed for the remaining images of the film to apply new black and white levels uniformly.

#### Quantification of receptor binding

On top of the hippocampal regions of interest (ROIs), four other ROIs were sampled bilaterally for their putative involvement in a corticolimbic circuit relevant to schizophrenia and hippocampal dysfunction (1): lower layers of the frontal regions (mPFC and ACC), dorsal striatum, ventral striatum and amygdala (Fig. S1). Non-specific binding for [$^{3}$H]-Ro15-4513 and [$^{3}$H]-flumazenil were negligible (see Fig. S2 for example image of non-specific binding).

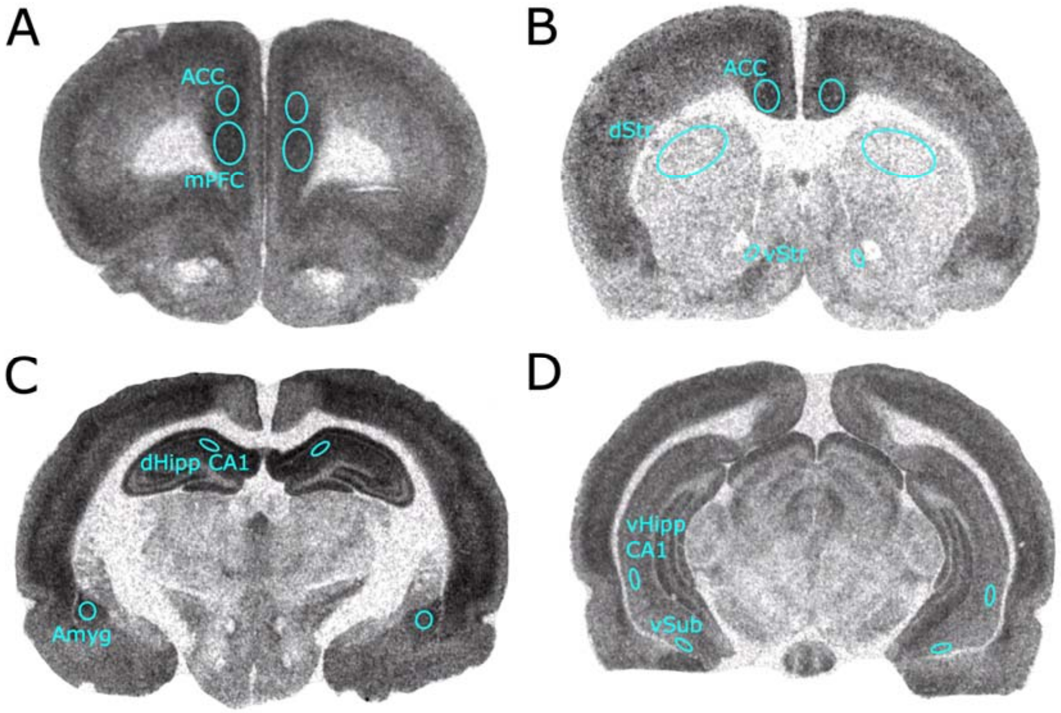


*Figure S1. All regions of interests sampled. A) lower layers of the medial prefrontal cortex (mPFC) and anterior cingulate cortex (ACC); B) ACC, dorsal Striatum (dStr), ventral Striatum (vStr); C) dorsal Hippocampus CA1 (dHipp CA1), amygdala (Amyg); D) ventral Hippocampus CA1 (vHipp CA1), ventral Subiculum (vSub).*

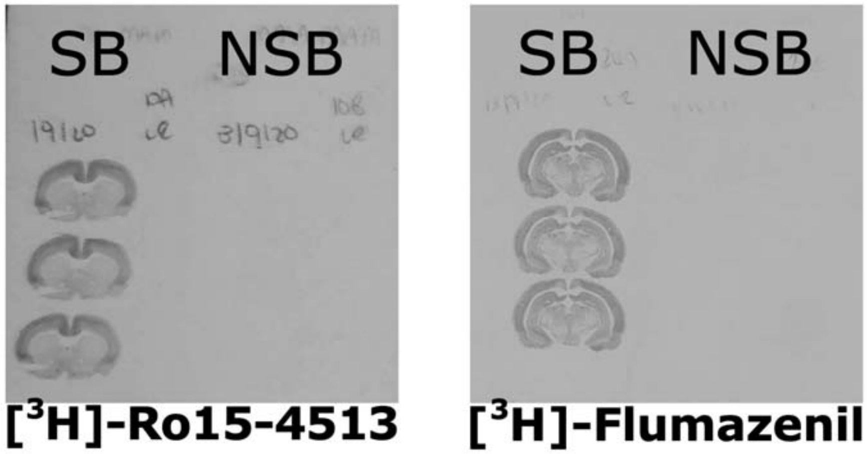


*Figure S2. Representative image of autoradiographs for [$^{3}$H]-Ro15-4513 and [$^{3}$H]-Flumazenil. Specific binding (SB) is displayed on the left of each image. Corresponding non-specific binding (NSB) is displayed on the right of each image. NSB was completely absent and therefore not measured.*

*Table S1. Final number of rats per experimental procedure*

| | EPM<br><br>n=71 | AIH<br><br>n=70 | [$^{3}$H]-Ro15-4513<br>n=62 | [$^{3}$H]-Flumazenil<br>n=62 | [$^{3}$H]-MK801<br><br>n=69 |
|---|---|---|---|---|---|
| SAL: VEH | 19 | 19 | 16 | 16 | 19 |
| SAL: DZP | 20 | 20 | 16 | 18 | 20 |
| MAM: VEH | 15 | 15 | 14 | 13 | 14 |
| MAM: DZP | 17 | 16 | 16 | 15 | 16 |

### Supplementary Results

### [$^3$H]-Ro15-4513

A 3-way mixed ANOVA (between subject: group (SAL/MAM), condition (VEH/DZP); repeated measures: ROIs) showed an overall effect of group indicating lower [$^3$H]-Ro15-4513 receptor binding ($F_{(1,58)}$ = 7.378, $p$ = 0.009) as well as an interaction effect (ROI x group, $F_{(6,341)}$ = 2.198, $p$ = 0.043). No main effect of condition was found ($F_{(1,58)}$ = 2.669, $p$ = 0.108). To investigate the interaction effect, *post hoc* tests corrected using the Benjamini Hochberg method was run following a 2-way mixed ANOVA where SAL:VEH and SAL:DZP were combined and MAM:VEH and MAM:DZP were combined into one group (between subject: group (SAL/MAM); repeated measures: ROIs). MAM rats exhibited decreases in [$^3$H]-Ro15-4513 binding in most regions ($q$ <0.05), but failed to reach significance in the amygdala and dorsal hippocampus CA1. $p$- and $q$-values for MAM versus saline effects are displayed in Table S1.

*Table S2. [$^3$H]-Ro15-4513 two-way ANOVA results per ROI*

| | Mean difference | *q* value | *p* value |
|---|---|---|---|
| mPFC+ACC | 5.175 | 0.032* | 0.023 |
| dStr | 0.9618 | 0.012* | 0.004 |
| vStr | 1.480 | 0.012* | 0.006 |
| Amyg | 2.092 | 0.116 | 0.116 |
| dHipp CA1 | 3.469 | 0.067^ | 0.057 |
| vHipp CA1 | 3.008 | 0.012* | 0.003 |
| vSub | 2.464 | 0.012* | 0.007 |

mPFC: medial prefrontal cortex; ACC: anterior cingulate cortex; dStr: dorsal striatum; vStr: ventral striatum; Amyg: amygdala; dHipp CA1: dorsal hippocampus CA1; vHipp CA1: ventral hippocampus CA1; vSub: ventral subiculum; ^$q$<0.1; *$q$<0.05; **$q$<0.01

#### [$^3$H]-flumazenil

Using a 3-way mixed ANOVA for [$^3$H]-flumazenil values, no MAM, Diazepam, or interaction effects were observed ($p$ > 0.05).

**[$^3$H]-MK801**

Lastly, a 3-way mixed ANOVA was conducted with [$^3$H]-MK801 values. A main effect of MAM indicating overall increased [$^3$H]-MK801 binding was observed ($F_{(1,67)}$ = 7.838, *p* = 0.007). An interaction effect failed to reach significance (ROI x MAM) ($F_{(6, 398)}$ = 2.023, *p* = 0.062), and no condition effect was found ($F_{(1,67)}$ = 0.040, *p* = 0.842).

### Behavior-autoradiography correlations

In addition to the correlation analyses outlined in the main text, receptor density values were correlated with behavior for any ROI that showed a significant main effect.

*Table S3. Behavior-autoradiography correlations*

| | | SAL:VEH | SAL:DZP | MAM:VEH | MAM:DZP | ALL RATS |
|---|---|---|---|---|---|---|
| Ro15 mPFC+ACC | AIH (totals) | -0.780 (<0.001)*** | -0.063 (0.823) | 0.004 (0.990) | -0.403 (0.122) | -0.373 (0.003)** |
| | EPM (Entries) | 0.097 (0.722) | -0.105 (0.700) | -0.450 (0.866) | 0.134 (0.621) | 0.188 (0.144) |
| | EPM (time) | -0.054 (0.843) | -0.119 (0.660) | -0.167 (0.567) | -0.059 (0.827) | 0.061 (0.638) |
| Ro15 dStr | AIH (totals) | -0.842 (<0.001)*** | 0.041 (0.884) | 0.037 (0.900) | -0.400 (0.175) | -0.334 (0.011)* |
| | EPM (Entries) | -0.091 (0.747) | -0.024 (0.929) | 0.028 (0.926) | -0.120 (0.695) | 0.138 (0.302) |
| | EPM (time) | -0.155 (0.582) | -0.043 (0.875) | 0.059 (0.842) | -0.284 (0.347) | 0.074 (0.582) |
| Ro15 vStr | AIH (totals) | -0.767 (0.001)** | 0.023 (0.934) | 0.097 (0.742) | -0.312 (0.258) | -0.317 (0.015)* |
| | EPM (Entries) | 0.069 (0.815) | -0.069 (0.801) | 0.080 (0.785) | 0.101 (0.722) | 0.207 (0.116) |
| | EPM (time) | -0.175 (0.551) | -0.058 (0.830) | 0.039 (0.896) | -0.074 (0.793) | 0.080 (0.549) |
| Ro15 dHipp CA1 | AIH (totals) | -0.709 (0.002)** | -0.140 (0.620) | 0.100 (0.733) | -0.277 (0.299) | -0.316 (0.013)* |
| | EPM (Entries) | 0.115 (0.671) | -0.095 (0.726) | 0.024 (0.936) | 0.193 (0.474) | 0.168 (0.191) |
| | EPM (time) | 0.062 (0.820) | -0.046 (0.865) | 0.021 (0.942) | -0.036 (0.895) | 0.106 (0.413) |
| Ro15 vHipp CA1 | AIH (totals) | -0.241 (0.368) | -0.140 (0.620) | -0.161 (0.582) | -0.455 (0.076) | -0.366 (0.004)** |
| | EPM (Entries) | -0.160 (0.554) | 0.351 (0.183) | 0.185 (0.526) | 0.088 (0.747) | 0.240 (0.061) |
| | EPM (time) | -0.021 (0.938) | 0.359 (0.172) | 0.369 (0.194) | -0.080 (0.768) | 0.263 (0.039)* |
| Ro15 vSub | AIH (totals) | -0.412 (0.113) | -0.264 (0.343) | -0.180 (0.538) | -0.384 (0.142) | -0.401 (0.001)** |
| | EPM (Entries) | -0.146 (0.591) | 0.161 (0.553) | 0.198 (0.498) | -0.002 (0.994) | 0.183 (0.154) |
| | EPM (time) | -0.018 (0.947) | 0.041 (0.880) | 0.451 (0.105) | -0.154 (0.570) | 0.175 (0.174) |
| MK801 mPFC+ACC | AIH (totals) | -0.237 (0.330) | 0.185 (0.510) | 0.198 (0.447) | 0.119 (0.326) | -0.237 (0.330) |
| | EPM (Entries) | 0.005 (0.983) | 0.158 (0.507) | -0.516 (0.049)* | 0.108 (0.679) | -0.047 (0.700) |
| | EPM (time) | -0.034 (0.891) | 0.269 (0.252) | -0.662 (0.007)** | 0.147 (0.575) | -0.033 (0.783) |
| MK801 dStr | AIH (totals) | 0.274 (0.256) | 0.110 (0.654) | -0.283 (0.307) | 0.203 (0.434) | 0.146 (0.228) |
| | EPM (Entries) | -0.020 (0.936) | 0.341 (0.142) | -0.398 (0.141) | -0.520 (0.032)* | -0.221 (0.064)^ |
| | EPM (time) | -0.016 (0.952) | 0.351 (0.130) | -0.247 (0.376) | -0.315 (0.218) | -0.111 (0.359) |
| MK801 vStr | AIH (totals) | 0.290 (0.228) | -0.005 (0.985) | -0.226 (0.437) | 0.456 (0.066)* | 0.174 (0.154) |
| | EPM (Entries) | 0.180 (0.461) | 0.193 (0.416) | -0.348 (0.223) | -0.455 (0.066)* | -0.1501 (0.215) |
| | EPM (time) | 0.193 (0.430) | 0.222 (0.348) | -0.404 (0.152) | -0.303 (0.238) | -0.1005 (0.408) |
| MK801 Amyg | AIH (totals) | 0.050 (0.839) | 0.360 (0.131) | -0.011 (0.970) | 0.499 (0.041)* | 0.280 (0.019)* |
| | EPM (Entries) | 0.133 (0.587) | 0.351 (0.129) | -0.503 (0.056)^ | 0.041 (0.874) | 0.005 (0.968) |
| | EPM (time) | 0.182 (0.457) | 0.283 (0.228) | -0.201 (0.472) | 0.123 (0.637) | 0.083 (0.492) |
| MK801 dHipp CA1 | AIH (totals) | 0.042 (0.864) | 0.264 (0.276) | 0.053 (0.851) | 0.655 (0.004)** | 0.318 (0.007)** |
| | EPM (Entries) | 0.061 (0.804) | 0.231 (0.327) | -0.565 (0.028)* | -0.242 (0.349) | -0.117 (0.332) |
| | EPM (time) | 0.135 (0.581) | 0.314 (0.178) | -0.576 (0.025)* | -0.269 (0.297) | -0.051 (0.673) |
| MK801 vHipp CA1 | AIH (totals) | -0.169 (0.489) | 0.356 (0.134) | 0.120 (0.671) | 0.341 (0.197) | 0.162 (0.183) |
| | EPM (Entries) | -0.309 (0.196) | 0.211 (0.372) | -0.542 (0.037)* | -0.326 (0.218) | -0.186 (0.123) |
| | EPM (time) | -0.330 (0.168) | 0.257 (0.274) | -0.484 (0.068)^ | -0.235 (0.381) | -0.131 (0.279) |
| MK801 vSub | AIH (totals) | -0.304 (0.206) | 0.321 (0.181) | -0.393 (0.165) | 0.275 (0.303) | 0.130 (0.292) |
| | EPM (Entries) | -0.303 (0.208) | 0.086 (0.720) | -0.090 (0.760) | 0.390 (0.136) | -0.122 (0.317) |
| | EPM (time) | -0.413 (0.079)^ | 0.051 (0.830) | -0.021 (0.944) | 0.312 (0.239) | -0.159 (0.192) |

mPFC: medial prefrontal cortex; ACC: anterior cingulate cortex; dStr: dorsal striatum; vStr: ventral striatum; Amyg: amygdala; dHipp CA1: dorsal hippocampus CA1; vHipp CA1: ventral hippocampus CA1; vSub: ventral subiculum.

Values are Pearson's *r* (*p*-value). ^$p<0.1$; *$p<0.05$; **$p<0.01$; ***$p<0.001$

### Supplementary References


1. Grace AA, Gomes FV (2019): The Circuitry of Dopamine System Regulation and its Disruption in Schizophrenia: Insights Into Treatment and Prevention. *Schizophrenia Bulletin*. 45:148-157.
